# Structural Homology and Electrostatic Potential Comparisons of Epitope Pair Candidates for Molecular Mimicry Triggering of Type 1 Diabetes Mellitus

**DOI:** 10.1101/2025.08.29.673145

**Authors:** Ryan Gardner, Joshua Wilkins, Sejal Mistry, Ramkiran Gouripeddi, Julio C. Facelli

**Affiliations:** Weber State University and University of Wisconsin-Madison; North Carolina A&T; Department of Biomedical Informatics; Utah Clinical and Translational Science Institute, The University of Utah

**Keywords:** Type 1 Diabetes Mellitus, Molecular Mimicry, Protein Structure Prediction

## Abstract

**Background:** Molecular mimicry, where foreign and self-peptides contain similar epitopes, can induce autoimmune responses. Identifying potential molecular mimics and studying their properties is key to understanding the onset of autoimmune diseases such as type 1 diabetes mellitus (T1DM). Previous work identified pairs of infectious epitopes (E_INF_) and T1DM epitopes (E_T1D_) that demonstrated sequence homology; however, structural homology was not considered. Correlating sequence homology with structural properties is important for translational investigation of potential molecular mimics. This work compares sequence homology with structural homology by calculating the structures and electrostatic potential surfaces of the epitope pairs identified in previous work from our laboratory.

**Results:** For each pair of E_INF_ and E_T1D_, the root mean square deviations (RMSD) were calculated between their predicted structures and their electrostatic potentials. Structures were predicted using the AlphaFold software program. Of the 53 epitope pairs considered here, only 10 did not exhibit any matching (i.e. less than 3 residues overlap). When considering all residues the RMSD ranges from 0.33 Å to 11.66 Å with an average of 2.68 Å. Twenty-two pairs (42%) have RMSD of less than 1.5 Å and 30 (58%) less than 3 Å.

**Conclusions:** Most of the E_INF_/E_T1D_ pairs selected by sequence homology show similar structural and electrostatic distributions, indicating that the E_INF_ may also bind to the same protein targets, i.e. the major histocompatibility complex molecules, for T1DM, leading to molecular mimicry onset of the disease. These findings suggest that searching for epitope pairs using sequence homology, a much less computationally demanding approach, leads to strong candidates for molecular mimicry that should be considered for further study. But structure homology, electrostatic potential calculations and full docking calculations may be necessary to advance the in-silico molecular mimicry predictions, which may be useful to select the most promising candidates for experimental studies.

## 1. Background

Molecular mimicry happens when external and self-peptides containing similar epitopes trigger a cross-reactive immune response (1-6). In susceptible individuals, molecular mimicry has been postulated as a mechanism for the onset of autoimmune diseases, including type 1 diabetes mellitus (T1DM) (7-9). T1DM is a chronic condition characterized by the patient inability to produce insulin due to an autoimmune insult. Insulin is a hormone produced in pancreatic β-cells that regulates carbohydrate metabolism. Understanding molecular mimicry can lead to the understanding of T1DM etiology and reveal novel paths for disease prevention and drug design.

The global incidence of T1DM has increased in recent years but its cause is largely unknown; however, it is thought that viral infections could be a significant contributor to this trend (10, 11). It is difficult to associate this steady rise of the incidence of T1DM with a single factor, such as genetics, given the heterogeneity of the condition (12), but it is likely the result of the complex interplay of multiple factors (i.e., genetic, hygiene, dietary, environmental, viral)., with molecular mimicry having an important role in this problem.

Previously, Mistry et al. (13) reported a pipeline for investigating molecular mimicry by finding homologous peptides implicated in T1DM etiology and infectious agents. This pipeline generates a prioritized list of infectious epitopes (E_INF_) showing sequence homology with T1DM epitopes (E_T1D_), with similar empirical binding affinity to specific major histocompatibility complexes (MHC), and similar empirical potential for T-cell immunogenicity. But the previous study did not consider structural homology of the epitope pairs (E_INF_/E_T1D_ pairs) found by sequence homology. While sequence homology may provide insight into potential molecular mimics, at a much lower computational cost, to further evaluate their potential it is necessary to compare structural characteristics. Furthermore, the electrostatics properties of the epitope pair must be compared because even when there is lack of structural similarity, the pair may show similar electrostatic potentials, revealing that the E_INF_ may still bind to the same protein targets (MHC molecules) and otherwise different electrostatic potential surfaces for structurally similar pairs may indicate lack of binding similarities. There are three advantages to a structural understanding of the epitope pairs. First, it provides insight into structural importance as proteins with similar structural homology have a great probability to share biochemical and physical properties. Secondly,, it may make possible the prediction of binding sites and other mechanisms resulting in molecular mimicry. Finally, it provides greater insight into drug development allowing for the disruption of cross-reactive immunity. Recent computational advancements make it possible the prediction of polypeptide structures and evaluate electrostatic properties. As shown in the results presented at a recent conference (14) it is now possible to use state of the art 3D protein structure prediction methods to study structural homology for relative large sets of epitope pairs. Moreover, in recent years there has been an increased interest in using silico methods to study antibody epitope interactions and their effect on immunity. For instance, Balbin et al. introduced Epitopedia, (5, 6) a comprehensive framework for studying molecular mimicry. Desta et al. (15) used in silico methods including structure prediction with Alphafold (16, 17) and PIPER docking (18) to model antibody-antigen interactions. Maleki et al. (19) used a comprehensive set of in silico tools to design a recombinant multi-epitope vaccine against influenza A, while Russo et al. used similar methods to design a COVID-19 vaccine (20).

Here we present a comprehensive structural study by comparing the predicted structures and their respective electrostatic potential of all the E_INF_/E_T1D_ pairs reported by Mistry et al. (13).

## 2. Results and Discussion

A summary of the results is presented in Table ST1. Table 1 presents the list of matching E_INF_/E_T1D_ pairs ordered by the RMSD values of the superposition of the maximum number of residues. Table ST2 presents the RMSD values of the superposition of best number of residues, (i.e. pruned overlap values). Table 2 presents the list nonmatching (i.e. overlap was observed for less than 3 amino acids) from the pair E_INF_/E_T1D_. In all tables the original amino acids are typed in red. The structural overlap of all E_INF_/E_T1D_ pairs considered here can be found in the deposit data structures at folder “Pairs Structure.zip” in the ZENODO repository: https://zenodo.org/records/11212016.

**Table 1:**
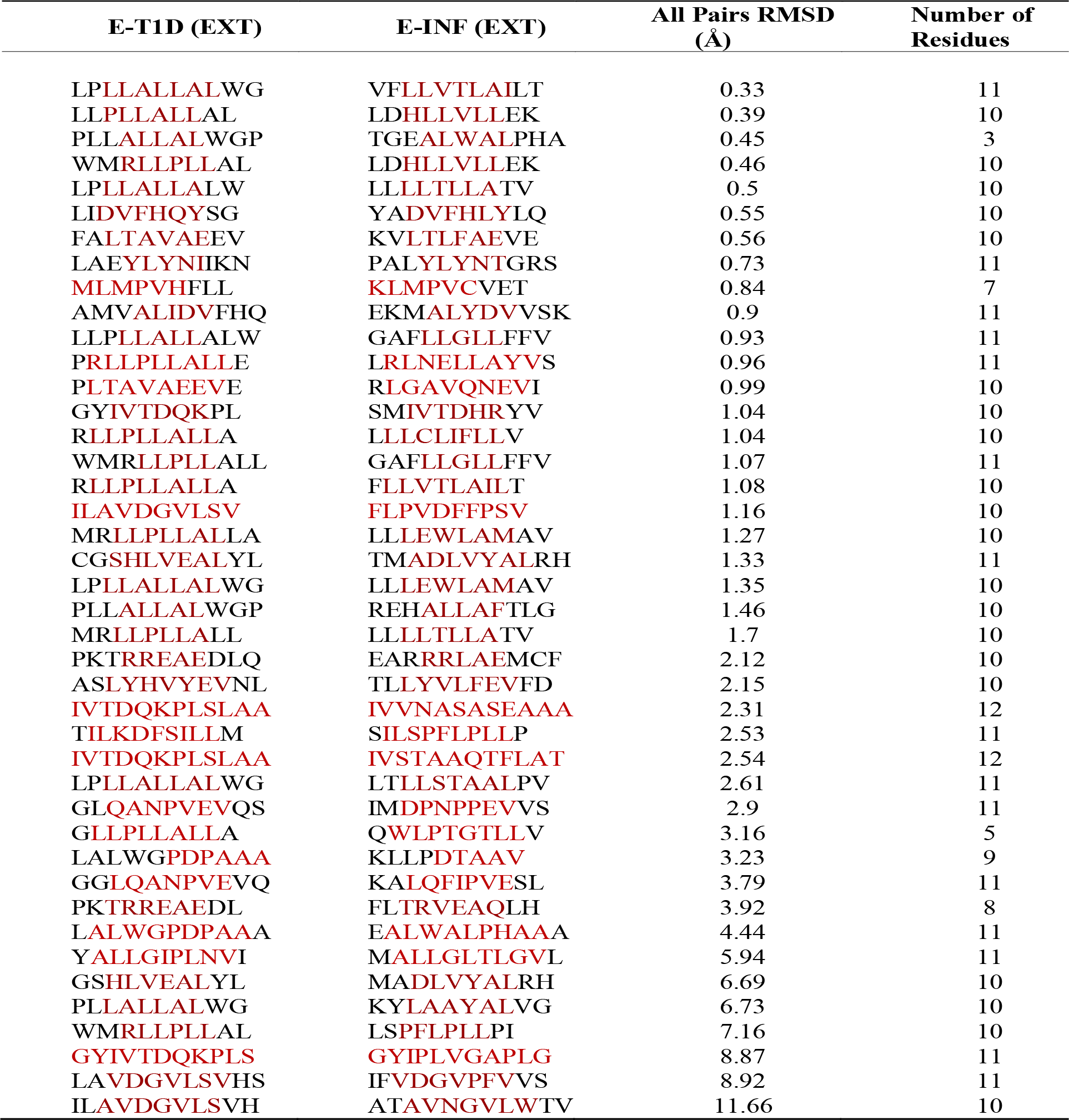
RMSD Between EINF/ET1D pairs for the Maximum of Overlapping Residues. Amino acids in red correspond to the original epitopes, those in black are added to reach the minimum required by AlphaFold.

**Table 2.**
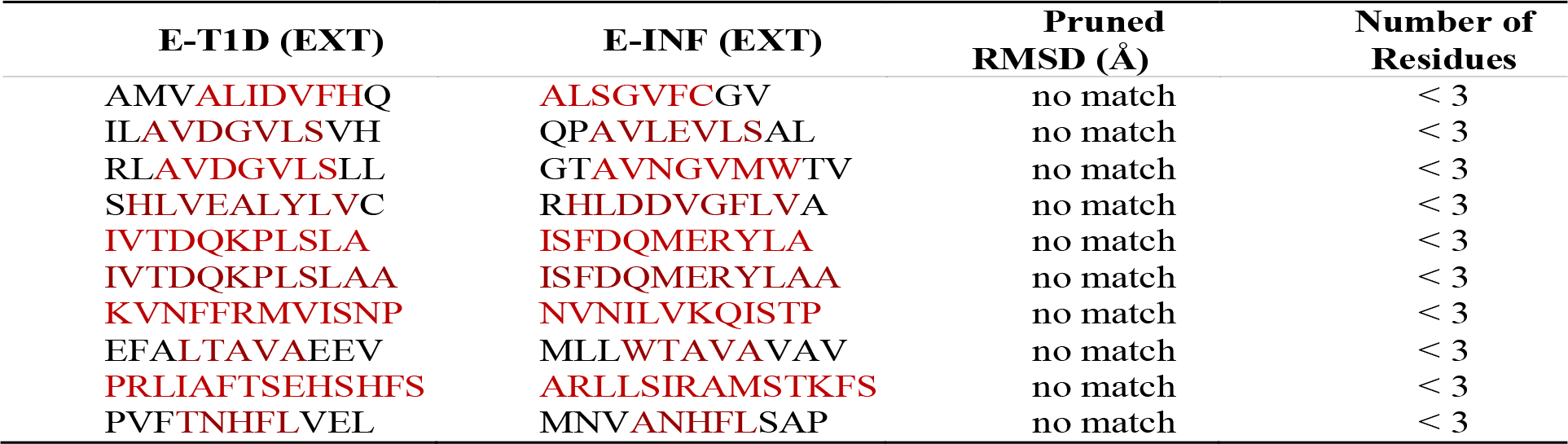
Nonmatching Epitopes Pairs. Amino acids in red correspond to the original epitopes, those in black are added to reach the minimum required by AlphaFold.

Of the 53 epitope pairs considered here, only 10 do not exhibit any matching (defined as less than 3 residues overlap). When considering all residues for the calculation of the RMSD, its range is from 0.33 Å to 11.66 Å with an average of 2.68 Å and having 22 pairs (42%) with RMSD of less than 1.5 Å and 30 (58%) less than 3 Å. It should be noted that the RMSD is not the only measure of quality presented in Table 1, as several epitope pairs in Table 1 show a low RMSD but for only a small number of residues, albeit larger than 3, but smaller than 10. Further examination of these pairs shows that they have very limited electrostatic similarity and likely are not good candidates for molecular mimicry (see Figure 1 and electrostatic surfaces in the folder “Pairs Structures.zip” in the data ZENODO repository: https://zenodo.org/records/11212016). The results for the pruned comparisons are presented in Table ST2, where the RMSD ranges from 0.09 Å to 96 Å, with 41 pairs (79%) having an RMSD of less than 1.5 Å. Moreover, it should be considered that the RMSD cutoff and the number of amino acids considered for homology is somehow arbitrary, because even epitopes with poor structural homology or homology restricted to a very small number of amino acids may still be effective in producing molecular mimicking in cases in which there is a very narrow biding site requiring only a small portion of the epitope to strongly bind. Only large-scale docking calculations can fully address this issue.

**Figure 1.**
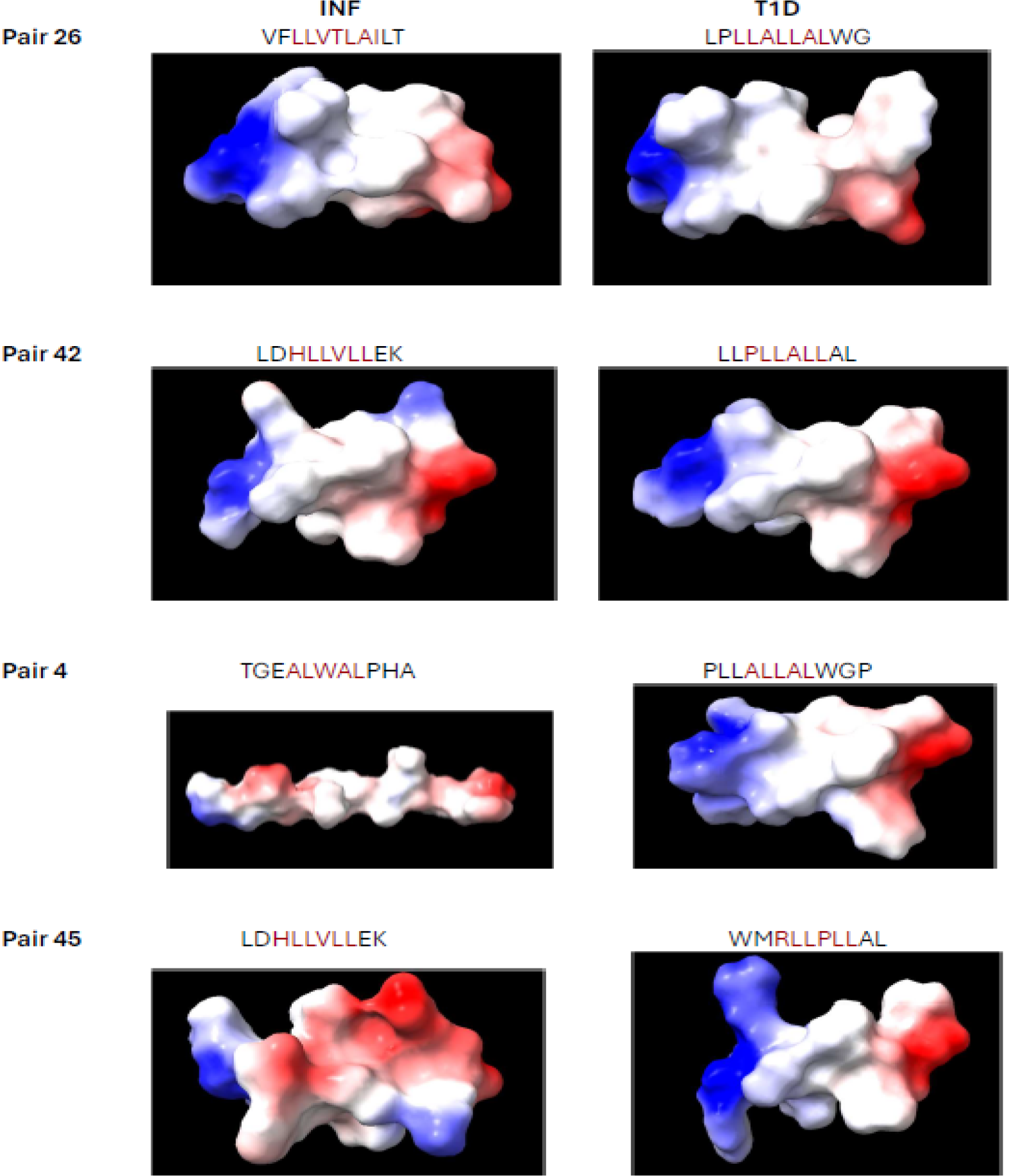
Comparison of the calculated electrostatic potential surfaces for all the best matching pairs from Table 1. The palette options were colored as follows: blue is defined to be negative; red is defined to be positive, and white is defined to be neutral. The nomenclature of the pair numbers can be found in Table S1 of the supplementary material.

The comparison of the calculated electrostatic potential surfaces for the best matching pairs from Table 1 is presented in Fig.1. For pairs 26, 42, and 45 we observe that they exhibit an excellent correlation in the electrostatic potential surfaces, with their corresponding small RMSDs, 0.33 Å, 0.39 Å and 0.46 Å, for 11, 10 and 10 residues, respectively. Of note, all pairs of the epitope’s secondary structure are the same for the E_INF_ and the E_T1D_ (all of them helix). The situation for pair #4, which exhibit and RMSD of 0.45 Å, but for only 3 residues, and where the E_INF_ shows a loop structure, while the E_T1D_ shows a helix folding is quite different, indicating than despite the low a RMSD this is not a good pair candidate for triggering T1DM by molecular mimicry. This is clearly consistent with the lack of similarity of the electrostatic potential surfaces depicted in Figure 1. This shows that small RMSD is a necessary condition, but not sufficient to identify good candidates for molecular mimicry. That a small RMSD is a necessary condition to find good candidates for E_INF_/E_T1D_ pairs likely to produce molecular mimicry, is further demonstrated in Figure SF1, where the calculated electrostatic potentials surface for all the nonmatching pairs from Table 2 are presented.

There are nine different antigens associated with the E_T1D_ studied here. Insulin shows five merged peptide domains, four of which are associated with HLA-A*02:01 and one is associated with HLA-B*08:01. GAD65 and IA-2 both contain three merged peptide domains associated with HLA-DRB4*01:01 or HLA-A*02:01. The remaining merged peptide domains were associated with HLA-A*02:01, with ZnT8 containing three merged peptide domains and S100B, KCNK16, PC2, ISL-1, and UCN3 all containing one merged peptide domain. The comparison of the structure and secondary folding of the E_INF_, E_T1D_ epitopes with the AlphaFold predicted structures of the corresponding amino acids sequences in the full antigen is presented in Figures 2-10.

**Figure 2.**
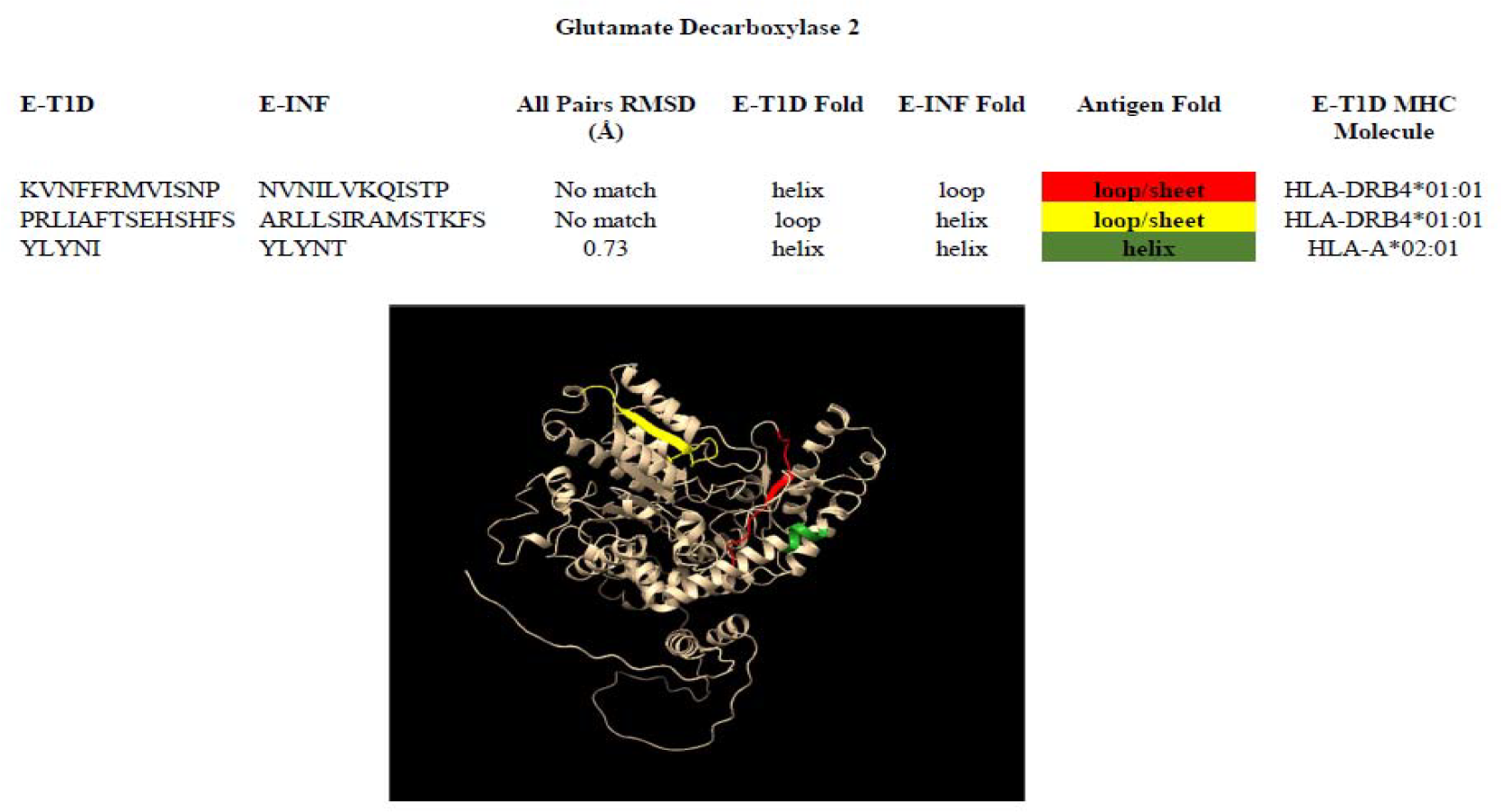
Infectious epitopes matching those from Glutamate Decarboxylase 2

Glutamate Decarboxylase 2 exhibits three regions in which epitopes can be found, they are located at positions Lys-553 to Pro-564 (red), Pro-271 to Ser-284 (yellow) and Tyr-480 to Ise-484 (green) as depicted in Fig. 2. The first two regions do not show a good match with the corresponding E_INF_ nor exhibit the same fold. The last region (green) shows a good match in both RMSD and similar fold than the YLYNT infectious epitope, which corresponds to the Human immunodeficiency virus type 1 (UniProt # P03369).

Insulin exhibits four regions in which epitopes can be found, they are Leu-7 to Leu-16 (red), Trp-17 to Ala-24 (orange), His-34 to Val-42 (yellow) and Thr-54 to Glu-60 (green) as depicted in Fig. 3. Within these regions there are 24 matching infections epitopes (by RMSD) and one not matching. From the 24 matching epitopes with small RMSD it is noted that three E_INF_, ALWAL, LAAYAL and WLPTGTLL show different fold than the E_T1D_ in either the free or in the antigen structures. Further examination of their electrostatic potential surfaces, see Fig. S2, shows that these are quite different indicating that most likely the corresponding epitope pairs do not have a high probability for molecular mimicry and therefore will not be considered strong molecular mimicry candidates. This leaves us with 21 strong candidates for molecular mimicry. The infectious epitopes, and their corresponding organisms and protein UniProt accession numbers, with a high likelihood to induce molecular mimicry are:

**Figure 3.**
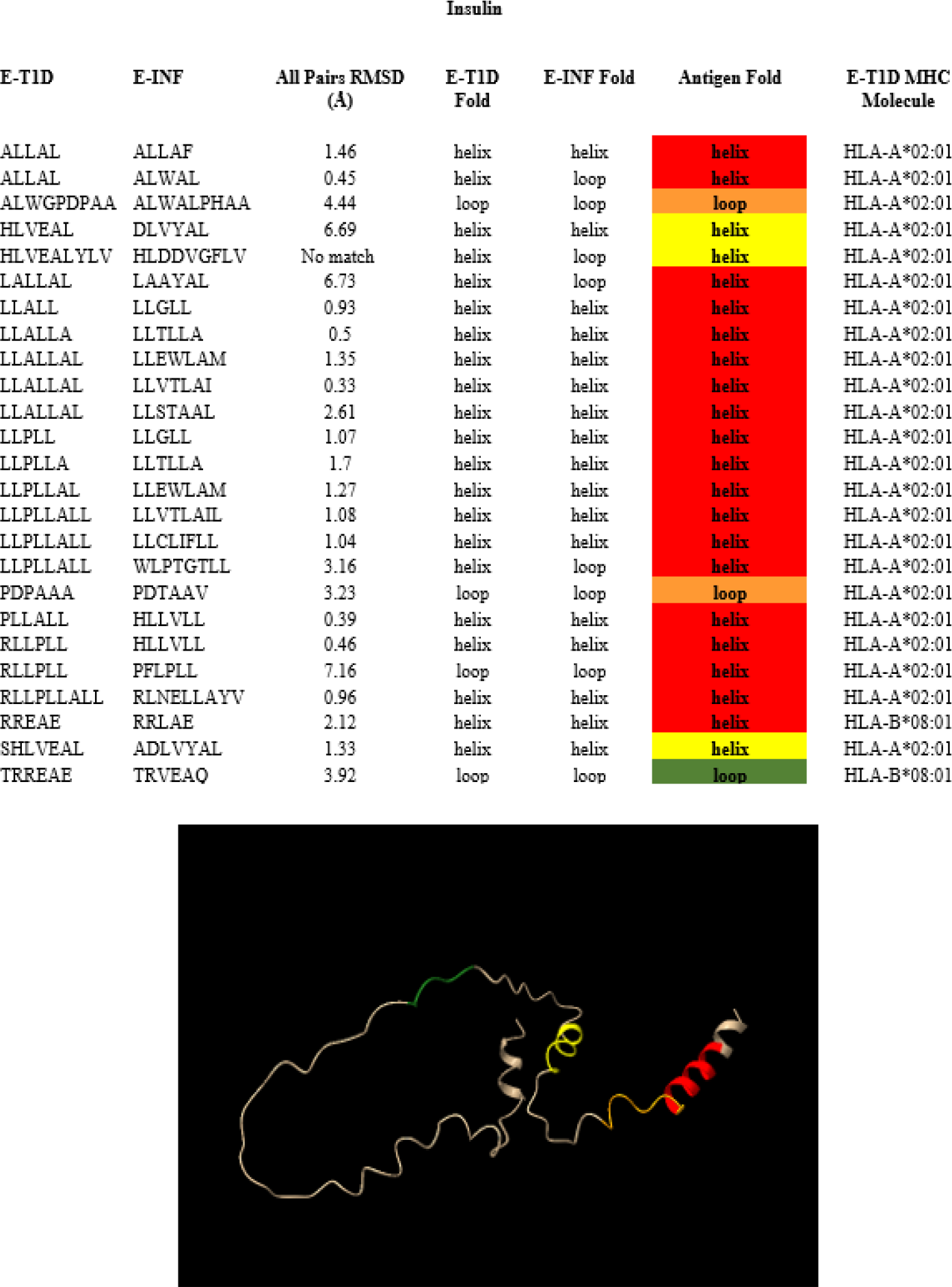
Infectious epitopes matching those from Insulin.

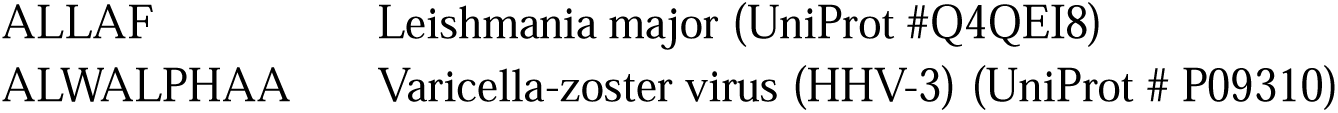
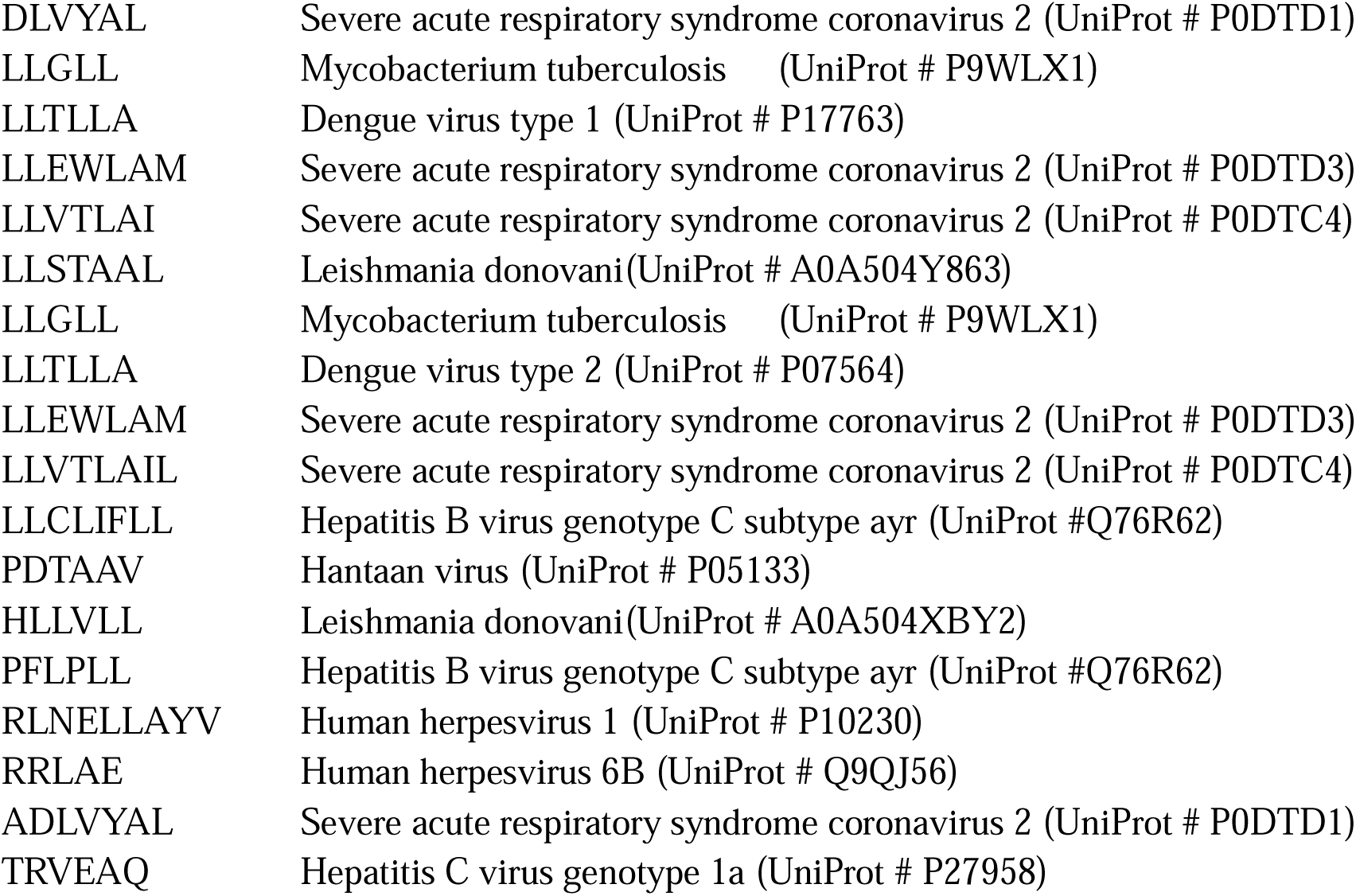

|  |  |
| --- | --- |
| ALLAF | Leishmania major (UniProt #Q4QEI8) |
| ALWALPHAA | Varicella-zoster virus (HHV-3) (UniProt # P09310) |
| DLVYAL | Severe acute respiratory syndrome coronavirus 2 (UniProt # P0DTD1) |
| LLGLL | Mycobacterium tuberculosis (UniProt # P9WLX1) |
| LLTLLA | Dengue virus type 1 (UniProt # P17763) |
| LLEWLAM | Severe acute respiratory syndrome coronavirus 2 (UniProt # P0DTD3) |
| LLVTLAI | Severe acute respiratory syndrome coronavirus 2 (UniProt # P0DTC4) |
| LLSTAAL | Leishmania donovani(UniProt # A0A504Y863) |
| LLGLL | Mycobacterium tuberculosis (UniProt # P9WLX1) |
| LLTLLA | Dengue virus type 2 (UniProt # P07564) |
| LLEWLAM | Severe acute respiratory syndrome coronavirus 2 (UniProt # P0DTD3) |
| LLVTLAIL | Severe acute respiratory syndrome coronavirus 2 (UniProt # P0DTC4) |
| LLCLIFLL | Hepatitis B virus genotype C subtype ayr (UniProt #Q76R62) |
| PDTAAV | Hantaan virus (UniProt # P05133) |
| HLLVLL | Leishmania donovani(UniProt # A0A504XBY2) |
| PFLPLL | Hepatitis B virus genotype C subtype ayr (UniProt #Q76R62) |
| RLNELLAYV | Human herpesvirus 1 (UniProt # P10230) |
| RRLAE | Human herpesvirus 6B (UniProt # Q9QJ56) |
| ADLVYAL | Severe acute respiratory syndrome coronavirus 2 (UniProt # P0DTD1) |
| TRVEAQ | Hepatitis C virus genotype 1a (UniProt # P27958) |

Insulin Gene Enhancer Protein ISL-1 exhibits one region in which epitopes can be found, it is Leu-277 to Val-284 (red) as depicted in Fig. 4. The two epitopes in this region show good match with the LQFIPVE and DPNPPEV infectious epitopes, which correspond to Hepatitis virus genotype C (UniProt # P27958) and Human immunodeficiency virus type 1 (UniProt # P03378), respectively.

**Figure 4.**
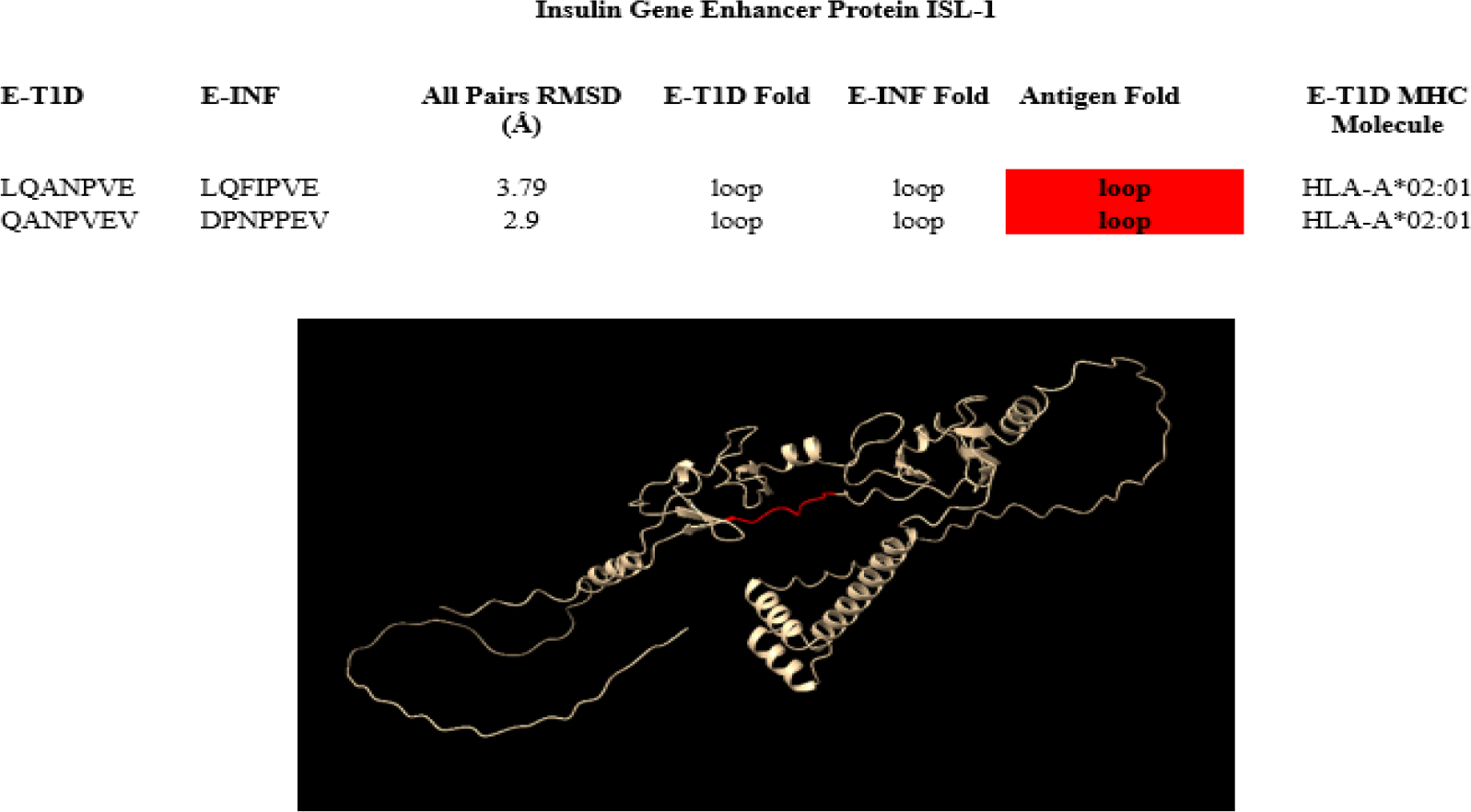
Infectious epitopes matching those from Insulin Gene Enhancer Protein ISL-1.

Neuroendocrine Convertase 2 exhibits one region in which epitopes can be found from Tyr-31 to Leu-35 (yellow) as depicted in Fig. 5. There is not a good matching infectious epitope for this antigen.

**Figure 5.**
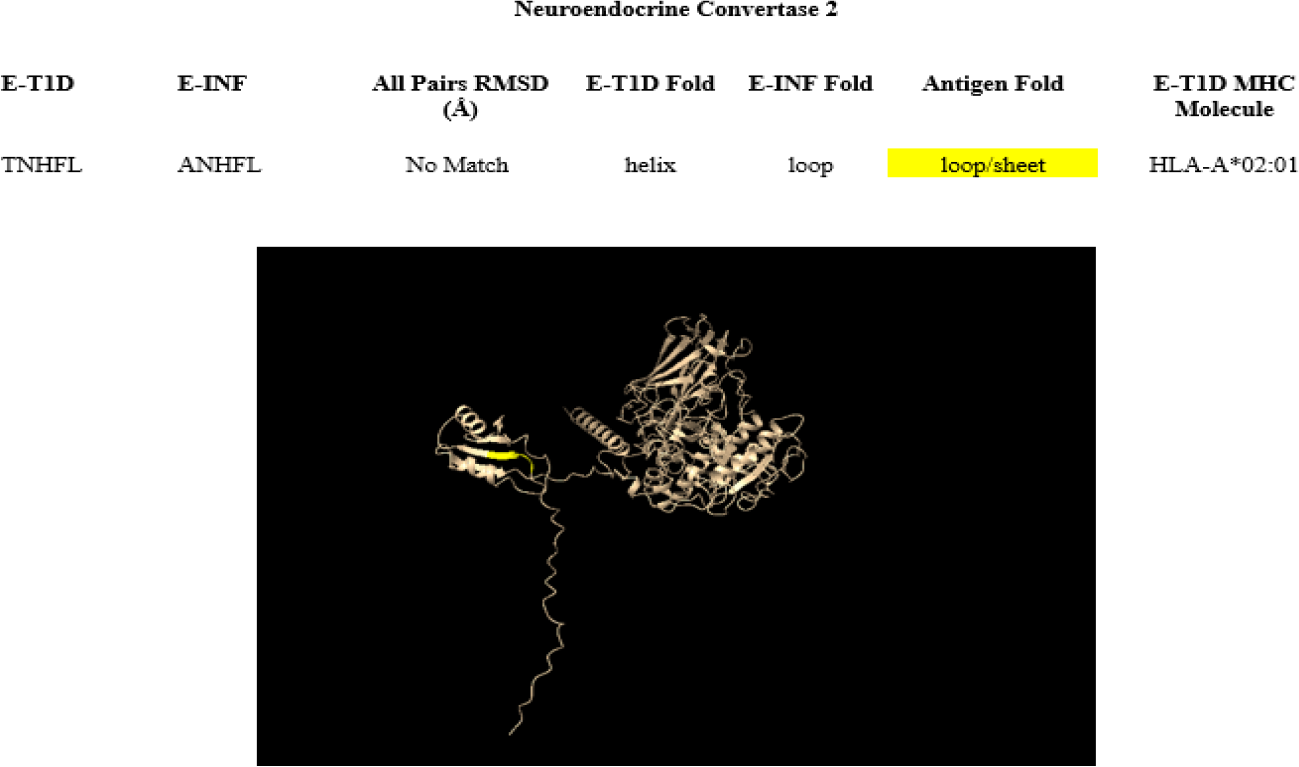
Infectious epitopes matching those from Neuroendocrine Convertase 2.

Potassium Channel Subfamily K Member 16 exhibits one region in which epitopes can be found at Ala-129 to Val-137 (green) as depicted in Fig. 6. There is a good match with the ALLGLTLGV infections epitope that corresponds to Human herpesvirus 1 (UniProt # P10211).

**Figure 6.**
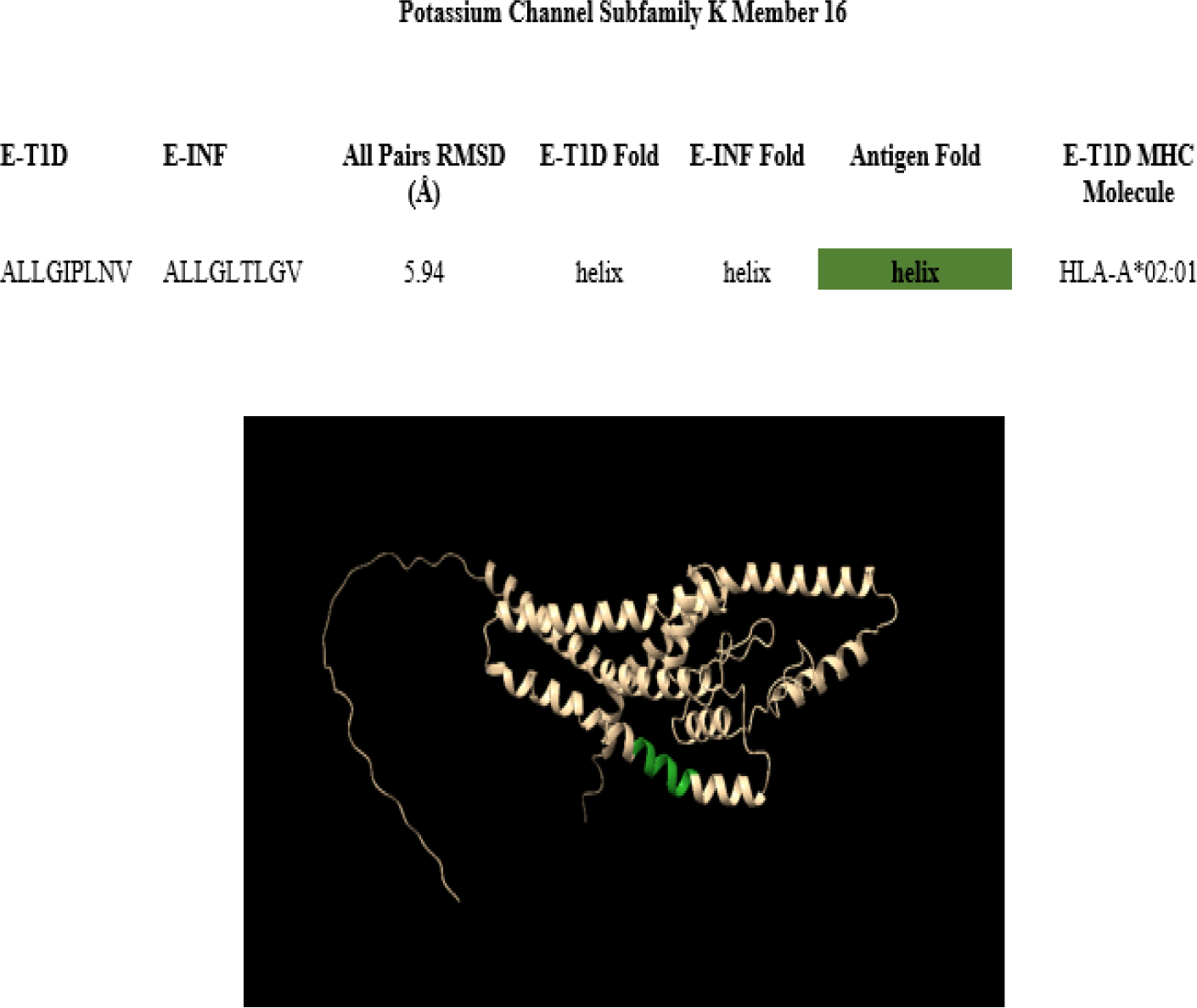
Infectious epitopes matching those from Potassium Channel Subfamily K Member 16.

Protein S100-B exhibits one region in which epitopes can be found at Ala-10 to Tyr-18 (red) as depicted in Fig. 7. There are two good matches with infectious epitopes in this region. They are ALIDV and DVFHQY, which correspond to Hepatitis C genotype 1a (UniProt # P27958) and Severe acute respiratory syndrome coronavirus 2 (UniProt # P0DTD1), respectively.

**Figure 7.**
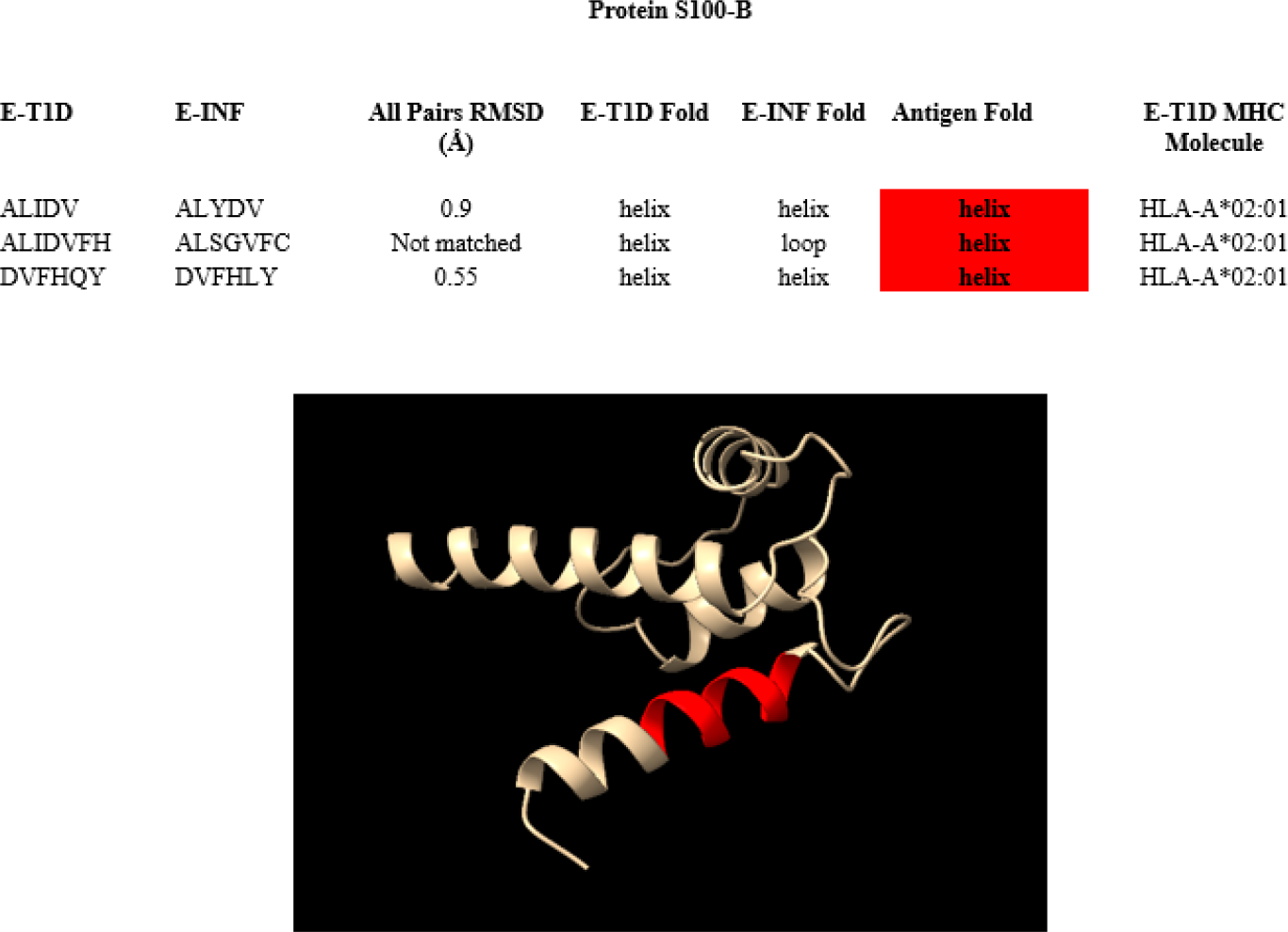
Infectious epitopes matching those from Protein S100-B.

Receptor-type tyrosine-protein phosphatase-like N exhibits three regions in which epitopes can be found at Gly-472 to Ala-485 (red), Leu-963 to Val-970 (green) and Leu-831 to Val-837 (yellow) as depicted in Fig. 8. In these regions we find seven matching infectious epitopes and three non-matching. From the seven matching epitopes with small RMSD it is noted that two E_INF_, LGAVQNEV and LYVLFEV show different folding than the E_T1D_ in either the free or in the antigen structures. Further examination of their electrostatic potential surfaces, see Fig. S2, shows that the electrostatic surfaces of these two pairs are quite similar, indicating that they are likely good candidates for molecular mimicry. The infectious epitopes, and their corresponding organisms and protein UniProt accession number, with a high likelihood to induce molecular mimicry are:

**Figure 8.**
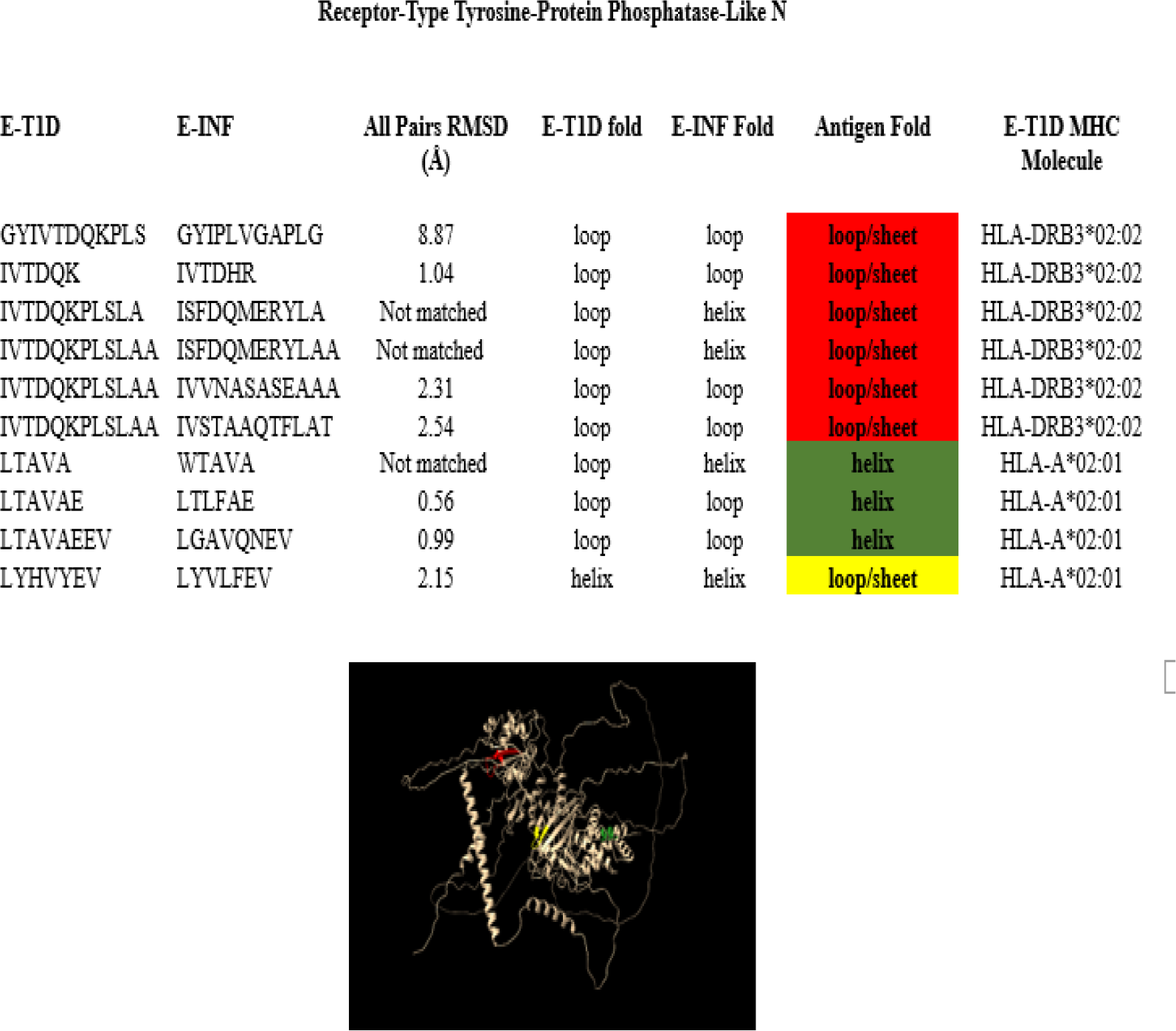
Infectious epitopes matching those from Receptor-type tyrosine-protein phosphatase-like N.

|  |  |
| --- | --- |
| GYIPLVGAPLG | Hepatitis C virus genotype 1a (UniProt # P27958) |
| IVTDHR | Mycobacterium tuberculosis (UniProt # P9WHH3) |
| IVVNASASEAAA | Coxiella burnetii (UniProt # Q83AF7) |
| IVSTAAQTFLAT | Hepatitis C virus genotype 1a (UniProt # P27958) |
| LTLFAE | Mycobacterium tuberculosis (UniProt # O53695) |
| LGAVQNEV | Hepatitis C virus genotype 1a (UniProt # P27958) |
| LYVLFEV | Human adenovirus C serotype 2 (UniProt # P03277) |

Urocortin-3 exhibits one region in which epitopes can be found at Met-1 to His-6 (yellow) as depicted in Fig. 9. Within this region we can find one matching infectious epitope, KLMPVC, which corresponds to severe acute respiratory syndrome coronavirus 2 (UniProt # P0DTD1).

**Figure 9.**
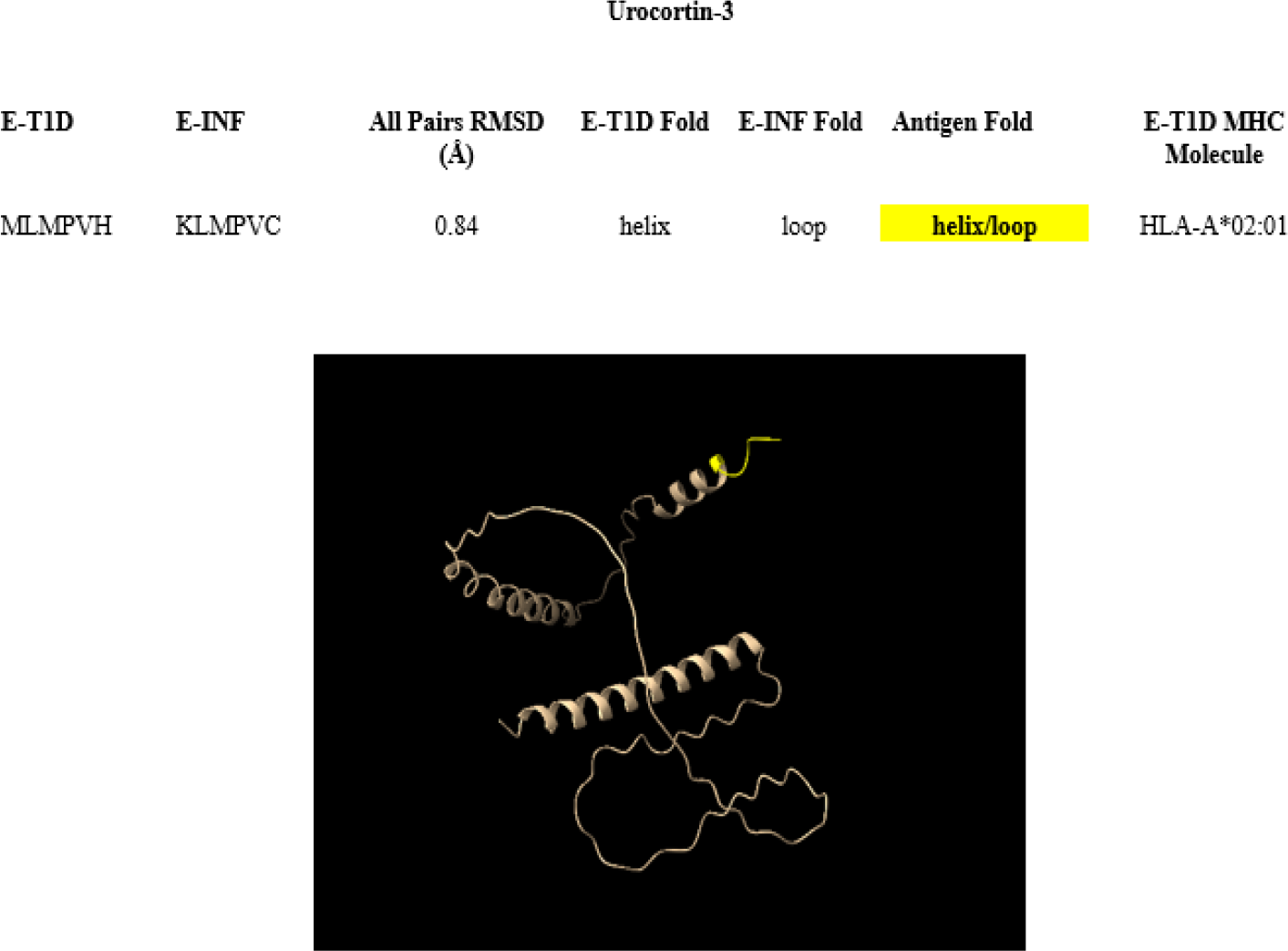
Infectious epitopes matching those from Urocortin-3.

Zinc Transporter 8 exhibits three regions in which epitopes can be found at Leu-107 to Leu-115 (yellow), Ile-266 to Leu-274 (red) and Ile-291 to Val-300 (green) depicted in Fig. 10. Within these regions we find four matching infectious epitopes with small RMSD but it is noted that the E_INF_ AVNGVLW shows a different fold than the corresponding E_T1D_ in either the free or in the antigen structures. Further examination of its electrostatic potential surface, see Fig. S2, shows that the electrostatic surface of the epitopes in this pair are quite different indicating that this is not a likely candidate for molecular mimicry. The infectious epitopes, and their corresponding organisms and proteins, with a high likelihood to induce molecular mimicry are:

**Figure 10.**
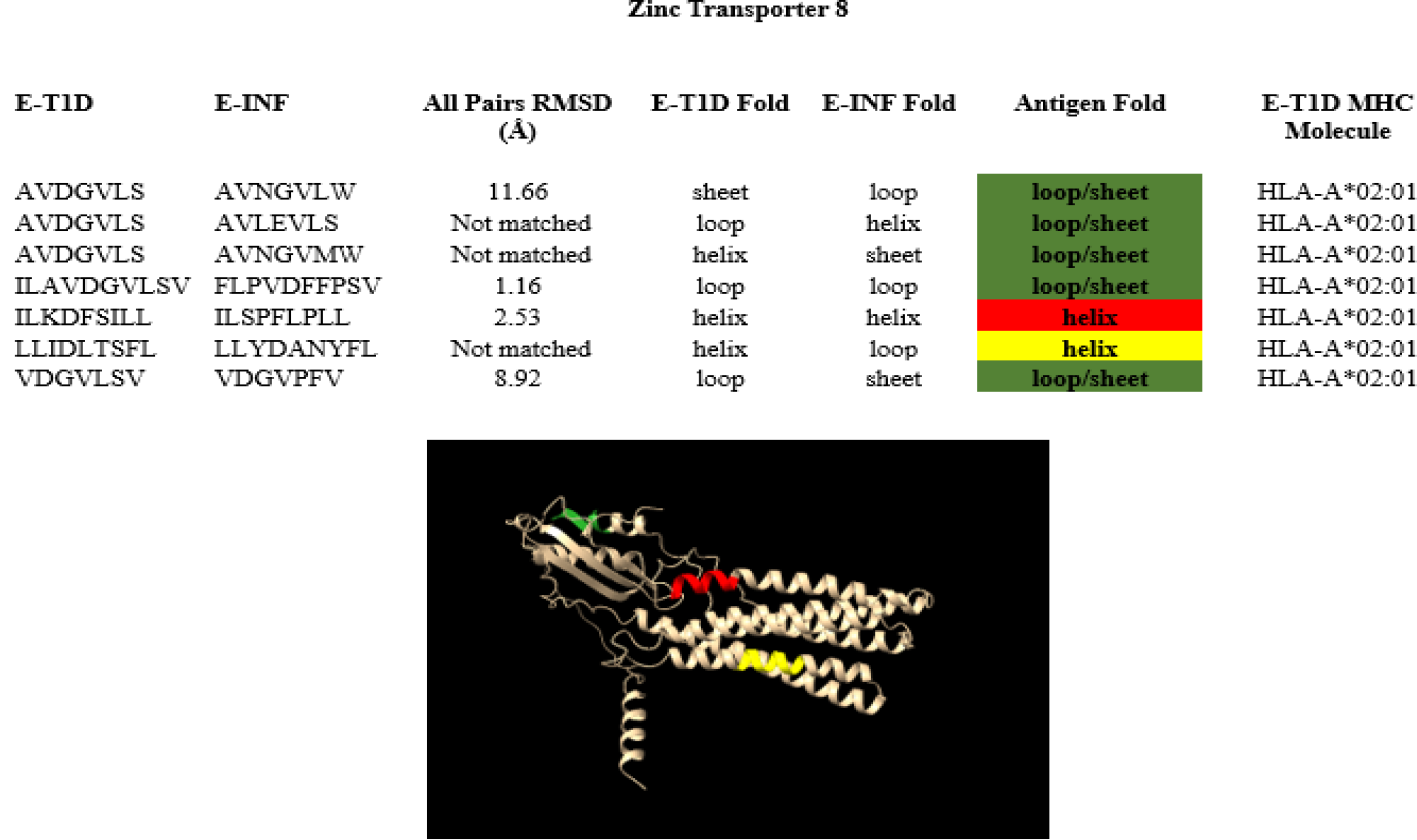
Infectious epitopes matching those from Zinc Transporter 8.

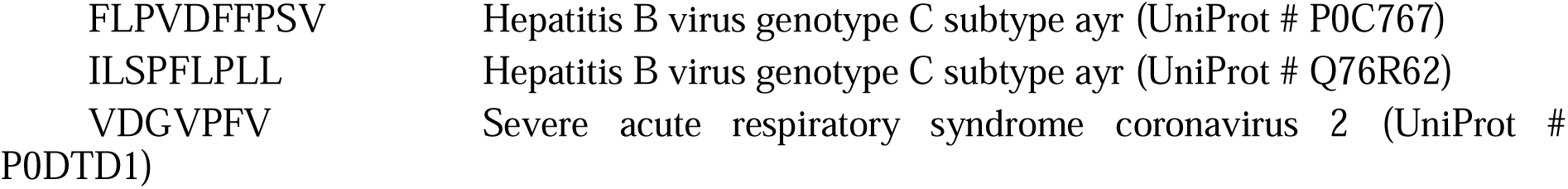

In total we have identified 23 different proteins in 15 different organisms that have a high likelihood of trigger molecular mimicry leading to the onset of T1DM. The list of organisms, protein UniProt reference number and the number of pairs in which an epitope of the protein has been found in this study are presented in Table 3. There are several organisms that have more than one protein capable of development of molecular mimicry, Hepatitis B (2), Human herpes virus 1 (2), Human immunodeficiency virus type 1 (2), Leishmania donovani (2), Mycobacterium tuberculosis (3) and severe acute respiratory syndrome coronavirus (3). It is quite notable that epitopes from proteins of the latest pathogen, commonly known as SARS-CoV-2 are present in nine E_INF/_E_T1D_ epitope pairs that have high likelihood to develop molecular mimicry. This is consistent with recent clinical observations that highlight the significant increase of T1DM cases after the COVID-19 pandemic (11), a topic of great interest that requires more study.

**Table 3.** List of organisms, their associated proteins and corresponding UniProt reference number, and the number of E_INF_/E_T1D_ pairs in which an epitope of the protein has been found in this study.

| Organism | UniProt Reference | Number of Epitope Paris |
| --- | --- | --- |
| Coxiella burnetii | Q83AF7 | 1 |
| Dengue virus type 1 (UniProt # P17763) | P17763 | 1 |
| Dengue virus type 2 | P07564 | 1 |
| Hantaan virus | P05133 | 1 |
| Hepatitis B virus genotype C subtype ayr | Q76R62 | 3 |
| Hepatitis B virus genotype C subtype ayr | P0C767 | 1 |
| Hepatitis C virus genotype 1a | P27958 | 6 |
| Human adenovirus C serotype 2 | P03277 | 1 |
| Human herpesvirus 1 | P10211 | 1 |
| Human herpesvirus 1 | P10230 | 1 |
| Human herpesvirus 6B | Q9QJ56 | 1 |
| Human immunodeficiency virus type 1 | P03369 | 1 |
| Human immunodeficiency virus type 1 | P03378 | 1 |
| Leishmania donovani | A0A504Y863 | 1 |
| Leishmania donovani | A0A504XBY2 | 1 |
| Leishmania major | Q4QEI8 | 1 |
| Mycobacterium tuberculosis | P9WLX1 | 2 |
| Mycobacterium tuberculosis | O53695 | 1 |
| Mycobacterium tuberculosis | P9WHH3 | 1 |
| Severe acute respiratory syndrome coronavirus 2 | P0DTD1 | 5 |
| Severe acute respiratory syndrome coronavirus 2 | P0DTC3 | 2 |
| Severe acute respiratory syndrome coronavirus 2 | P0DTD4 | 2 |
| Varicella-zoster virus (HHV-3) | P09310 | 1 |

In a recent paper Mistry et al. (24) identified sequences of infectious conditions leading to an increased likelihood of the onset of T1DM. The study used the TEDDY data set (25) version 25, and most of the conditions studied there cannot be directly associated with an specific pathogen. Exceptions are varicella and sixth disease, whose pathogens are also found in Table 3. Several pathogens in Table 3, like Dengue, Leishmania, tuberculosis, etc., are not likely to be found in the TEDDY study as these diseases are uncommon in the TEDDY population. Finally, it should be noted that version 25 of the TEDDY study predates the COVID-19 pandemic and therefore we did not expect to find any information on the importance of the SARS-CoV-2 infections in the study reported in reference (19).

## 3. Methods

Previous work (13) identified, catalogued, and indexed 53 unique pairs of infectious epitopes (E_INF_) and T1DM epitopes (E_T1D_) that demonstrated sequence homology and potential for molecular mimicry. Here, we predicted the structure of all these epitopes using AlphaFold (16, 17) version 2.3.2, with default parameters. AlphaFold is a new machine learning approach that incorporates previous knowledge about protein structure and that leveraging multi-sequence alignments into the design of a deep learning algorithm can predict protein structures with great accuracy. Because this method requires a sequence of at least ten amino acids and many of the epitope pairs did not meet this requirement, we expanded the epitopes using the full protein sequences from UniProt (https://www.uniprot.org/). Both the E_INF_ and E_T1D_ sequences were expanded symmetrically at both ends to reach the minimum length required by AlphaFold. The expansion of the epitopes to the minimum required length is a limitation in our approach, but it should be considered as an approximation necessary when using any state-of-the-art 3D protein structure prediction method (14). While in absence of experimental evidence it is difficult to ascertain the exact implications of this approximation, we compared the structures of the best matching epitopes, LP**LLALLAL**WG and VF**LLVTLAI**LT, with those extended even more by adding three additional amino acids at each end, LALLP**LLALLAL**WGRRN and AFVV**FLLVTLAI**LTALR. The AlphaFold predicted structures of these four epitopes show no differences in the central regions corresponding to the original ones. The best match of the central regions of LP**LLALLAL**WG and VF**LLVTLAI**LT shows an RMSD of 0.33 Å over 11 pairs, that can be compared with the overall superposition of all the structures including the ones expanded by additional six amino acids, which show an RMSF of 0.30 Å over 11 pairs. The superposition, not shown here, shows almost perfect overlap in the original epitopes region, supporting the use of expansion of the epitopes as done here and arguing that end effects may not affect the results presented here.

All sequences of the epitopes studied here are in the supplementary material (Complete Results, Table ST1), where the original amino acids are typed in red. The antigen structures were also obtained with AlphaFold from Ref. (16). Structures were compared, visualized with UCSF ChimeraX (21-23), using default parameters. The RMSD was calculated for each E_INF_/E_T1D_ pair by utilizing the “matchmaker” tool in ChimeraX and the electrostatic potential surfaces of each epitope was calculated also using ChimeraX. In ChimraX, the calculation of the electrostatic potentials is done by the Adaptive Poisson-Boltzmann Solver, which generates a grid-based potential map that ChimeraX visually represents on a molecular surface. The palette options were colored as follows: blue is defined to be negative; red is defined to be positive, and white is defined to be neutral. All the results presented here are based on the best ranked structure generated by AlphaFold. Full results of the AlphaFold calculation are in file “AF_2.3.2 Epitopes.zip” in the ZENODO repository: https://zenodo.org/records/11212016.

## 4. Conclusions

The results presented here show that the Mistry et al.’ sequence homology pipeline (13) is successful for finding epitope pairs that also show structural homology and in finding epitope pairs that are good candidates for triggering molecular mimicry, making them targets for further investigation. The use of this pipeline allowed us to perform an insightful investigation of structural homology and the electrostatic potential surfaces similarities of the Mistry’s epitope pairs, thereby avoiding the more computationally demanding method of structural screening all possible pairs of epitopes. The results also provide evidence that sequence and structural homology are not the whole picture, and that electrostatic potential and folding should be considered. Still full docking calculations may be necessary to advance the in-silico molecular mimicry predictions.

Future work including the investigation of the unmatched pairs and documenting more details of the structural outliers is needed to determine binding properties using molecular docking. This may provide better insights into the nature of molecular mimicry leading to the onset of T1DM.

## Supporting information

All suplementary material

## Author Contributions

RG and JW conceptualized the research, performed the calculations, and prepared the initial report. SM conceptualized the research and contributed to the writing of the manuscript. RG SM conceptualized the research and contributed to the writing of the manuscript. JCF SM conceptualized the research, performed the final calculations, provided overall supervision of the project, and contributed to the writing of the manuscript. All the authors reviewed and approved the final version of the manuscript.

## Data Availability Statement

The data presented in this study are openly available in ZENDO: https://zenodo.org/records/11212016.

## References

1. Cusick MF, Libbey JE, Fujinami RS. Molecular mimicry as a mechanism of autoimmune disease. Clin Rev Allergy Immunol. 2012;42(1):102–11.

2. Rojas M, Restrepo-Jiménez P, Monsalve DM, Pacheco Y, Acosta-Ampudia Y, Ramírez-Santana C, et al. Molecular mimicry and autoimmunity. Journal of Autoimmunity. 2018;95:100–23.

3. Blank M, Barzilai O, Shoenfeld Y. Molecular mimicry and auto-immunity. Clin Rev Allergy Immunol. 2007;32(1):111–8.

4. Fujinami RS, von Herrath MG, Christen U, Whitton JL. Molecular mimicry, bystander activation, or viral persistence: infections and autoimmune disease. Clin Microbiol Rev. 2006;19(1):80–94.

5. Balbin CA, Nunez-Castilla J, Stebliankin V, Baral P, Sobhan M, Cickovski T, et al. Epitopedia: identifying molecular mimicry between pathogens and known immune epitopes. ImmunoInformatics. 2023;9:100023.

6. Stebliankin V, Baral P, Balbin C, Nunez-Castilla J, Sobhan M, Cickovski T, et al. EMoMiS: A Pipeline for Epitope-based Molecular Mimicry Search in Protein Structures with Applications to SARS-CoV-2. bioRxiv; 2022.

7. Roep BO, Hiemstra HS, Schloot NC, De Vries RR, Chaudhuri A, Behan PO, et al. Molecular mimicry in type 1 diabetes: immune cross-reactivity between islet autoantigen and human cytomegalovirus but not Coxsackie virus. Ann N Y Acad Sci. 2002;958:163–5.

8. Honeyman MC, Stone NL, Harrison LC. T-cell epitopes in type 1 diabetes autoantigen tyrosine phosphatase IA-2: potential for mimicry with rotavirus and other environmental agents. Mol Med. 1998;4(4):231–9.

9. Christen U, Bender C, von Herrath MG. Infection as a cause of type 1 diabetes? Curr Opin Rheumatol. 2012;24(4):417–23.

10. Gale EAM. The Rise of Childhood Type 1 Diabetes in the 20th Century. Diabetes. 2002;51(12):3353–61.

11. Kamrath C, Holl RW, Rosenbauer J. Elucidating the Underlying Mechanisms of the Marked Increase in Childhood Type 1 Diabetes During the COVID-19 Pandemic—The Diabetes Pandemic. JAMA Network Open. 2023;6(6):e2321231–e.

12. Ogrotis I, Koufakis T, Kotsa K. Changes in the Global Epidemiology of Type 1 Diabetes in an Evolving Landscape of Environmental Factors: Causes, Challenges, and Opportunities. Medicina (Kaunas). 2023;59(4).

13. Mistry S, Gouripeddi R, Facelli JC. Prioritization of infectious epitopes for translational investigation in type 1 diabetes etiology. Journal of Autoimmunity. 2023;140:103115.

14. Gardner R, Wilkins J, Mistry S, Gouripeddi R, Facelli JC, editors. Structural Homology of Epitope Binding Mimicry in the Onset of Type 1 Diabetes Mellitus. 2023 IEEE International Conference on Bioinformatics and Biomedicine (BIBM); 2023: IEEE.

15. Desta IT, Kotelnikov S, Jones G, Ghani U, Abyzov M, Kholodov Y, et al. Mapping of antibody epitopes based on docking and homology modeling. Proteins. 2023;91(2):171–82.

16. Jumper J, Evans R, Pritzel A, Green T, Figurnov M, Ronneberger O, et al. Highly accurate protein structure prediction with AlphaFold. Nature. 2021;596(7873):583–9.

17. Varadi M, Anyango S, Deshpande M, Nair S, Natassia C, Yordanova G, et al. AlphaFold Protein Structure Database: massively expanding the structural coverage of protein-sequence space with high-accuracy models. Nucleic Acids Research. 2022;50(D1):D439–D44.

18. Kozakov D, Brenke R, Comeau SR, Vajda S. PIPER: an FFT-based protein docking program with pairwise potentials. Proteins. 2006;65(2):392–406.

19. Maleki A, Russo G, Parasiliti Palumbo GA, Pappalardo F. In silico design of recombinant multi-epitope vaccine against influenza A virus. BMC Bioinformatics. 2022;22(14):617.

20. Russo G, Pennisi M, Fichera E, Motta S, Raciti G, Viceconti M, et al. In silico trial to test COVID-19 candidate vaccines: a case study with UISS platform. BMC Bioinformatics. 2020;21(Suppl 17):527.

21. Pettersen EF, Goddard TD, Huang CC, Meng EC, Couch GS, Croll TI, et al. UCSF ChimeraX: Structure visualization for researchers, educators, and developers. Protein Sci. 2021;30(1):70–82.

22. Goddard TD, Huang CC, Meng EC, Pettersen EF, Couch GS, Morris JH, et al. UCSF ChimeraX: Meeting modern challenges in visualization and analysis. Protein Sci. 2018;27(1):14–25.

23. Pettersen EF, Goddard TD, Huang CC, Couch GS, Greenblatt DM, Meng EC, et al. UCSF Chimera—A visualization system for exploratory research and analysis. Journal of Computational Chemistry. 2004;25:1605–12.

24. Mistry S, Gouripeddi R, Raman V, Facelli JC. Sequential data mining of infection patterns as predictors for onset of type 1 diabetes in genetically at-risk individuals. Journal of Biomedical Informatics. 2023:104385.

25. Lönnrot M, Lynch K, Larsson HE, Lernmark Å, Rewers M, Hagopian W, et al. A method for reporting and classifying acute infectious diseases in a prospective study of young children: TEDDY. BMC Pediatrics. 2015;15(1):24.

