## Supplementary material for "Structural Homology and Electrostatic Potential Comparisons of Epitope Pair Candidates for Molecular Mimicry Triggering of Type 1 Diabetes Mellitus": All suplementary material: Figure SF1.pdf

| | INF | T1D |
|---|---|---|
| **Pair 2** | ALSGVFCGV | AMVALIDVFHQ |
| | 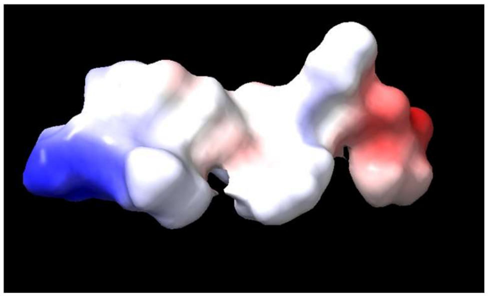 | 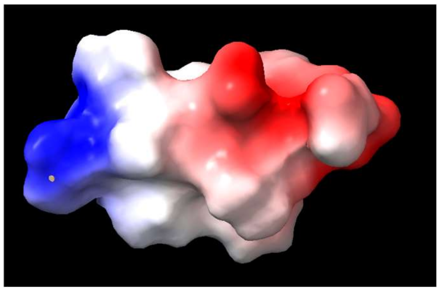 |
| **Pair 8** | QPAVLEVLSAL | ILAVDGVLSVH |
| | 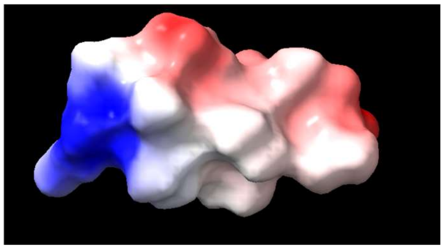 | 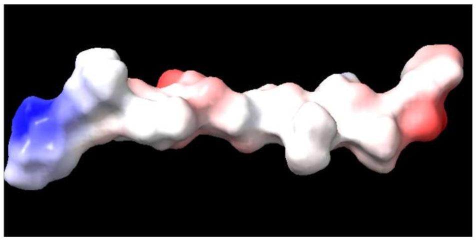 |
| **Pair 9** | GTAVNGVMWTV | RLAVDGVLSLL |
| | 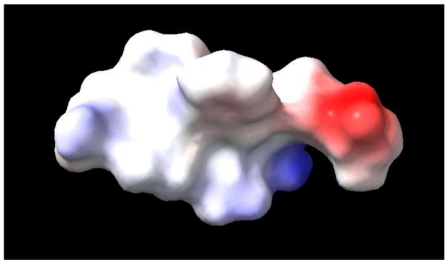 | 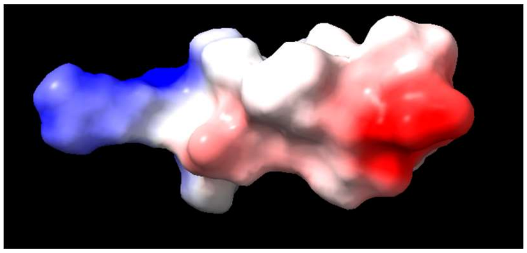 |
| **Pair 13** | RHLDDVGFLVA | SHLVEALYLVC |
| | 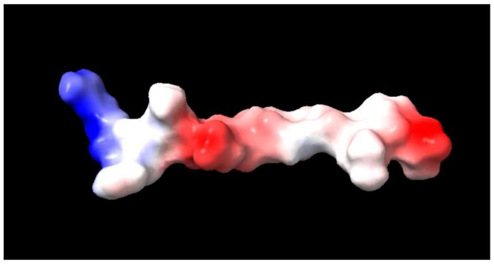 | 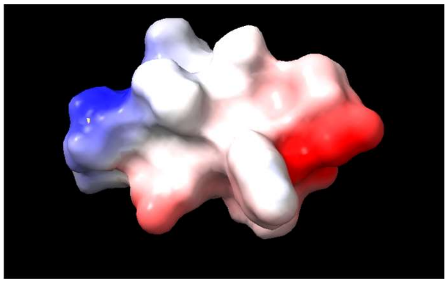 |

**Pair 17** ISFDQMERYLA IVTDQKPLSLA

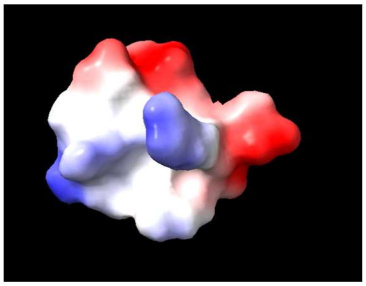

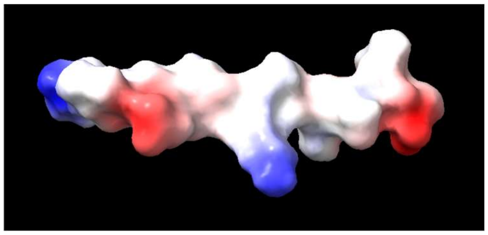

**Pair 18** ISFDQMERYLAA IVTDQKPLSLAA

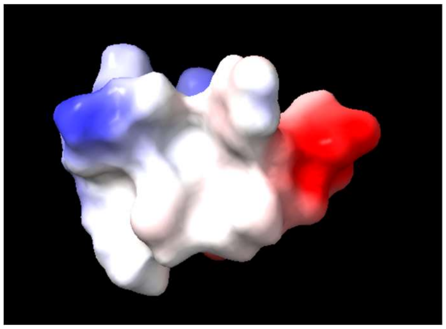

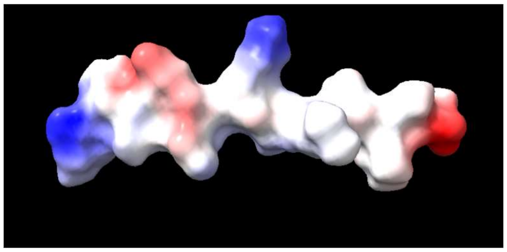

**Pair 21** NVNILVKQISTP KVNFFRMVISNP

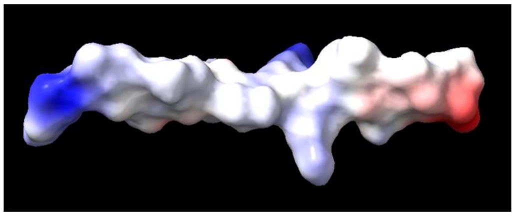

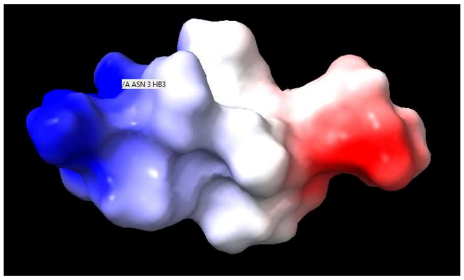

**Pair 36** MLLWTAVAVAV EFALTAVAEEV

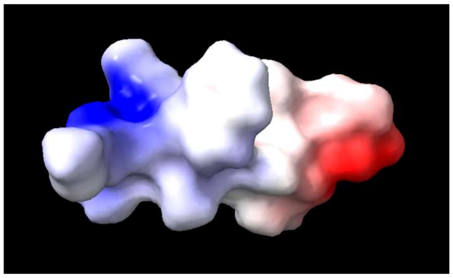

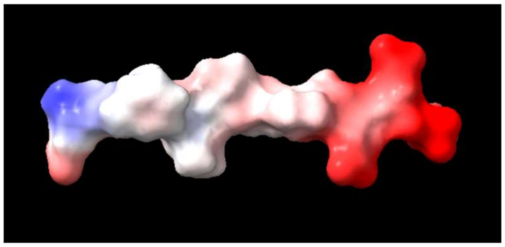

**Pair 43** ARLLSIRAMSTKFS PRLIAFTSEHSHFS

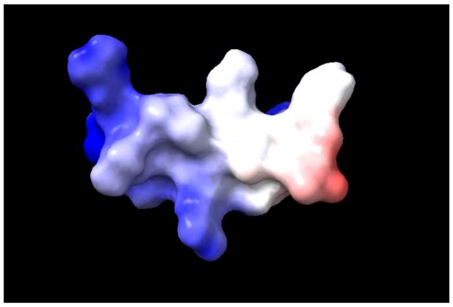

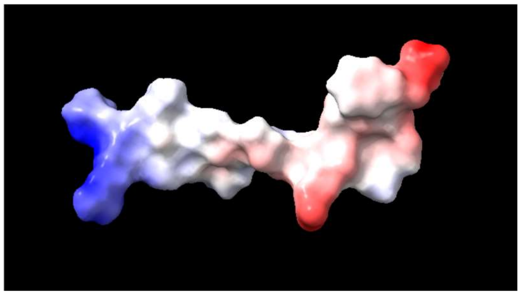

**Pair 50** MNVANHFLSAP PVFTNHFLVEL

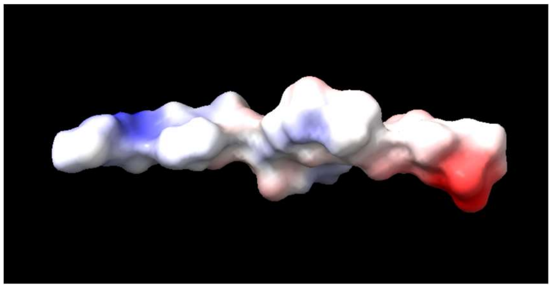

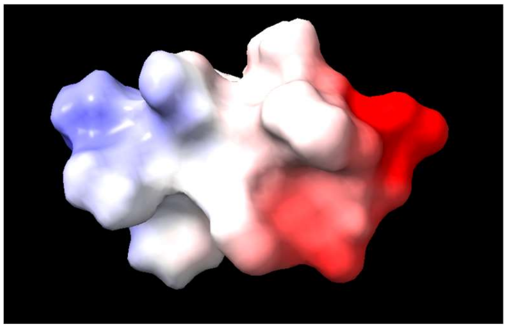
