## Supplementary material for "Structural Homology and Electrostatic Potential Comparisons of Epitope Pair Candidates for Molecular Mimicry Triggering of Type 1 Diabetes Mellitus": All suplementary material: Figure SF2.pdf

| | INF | T1D |
|---|---|---|
| **Pair 4** | **ALWAL** | **ALLAL** |
| | 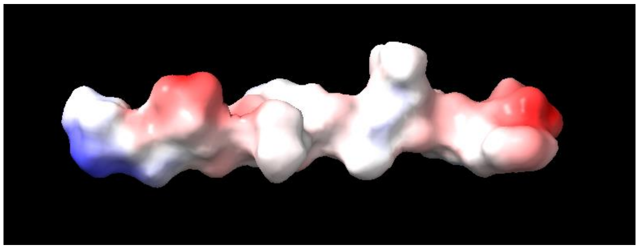 | 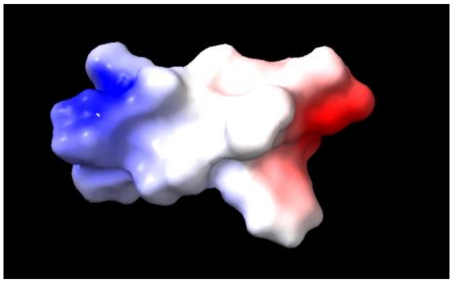 |
| **Pair 22** | **LAAYAL** | **LALLAL** |
| | 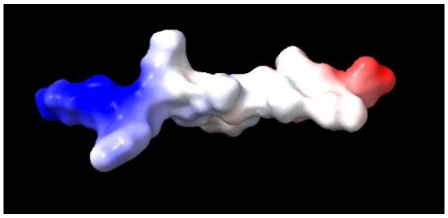 | 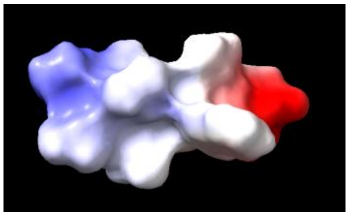 |
| **Pair 34** | **WLPTGTLL** | **LLPLLALL** |
| | 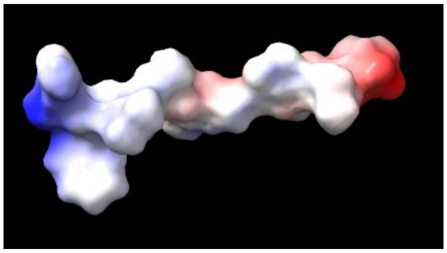 | 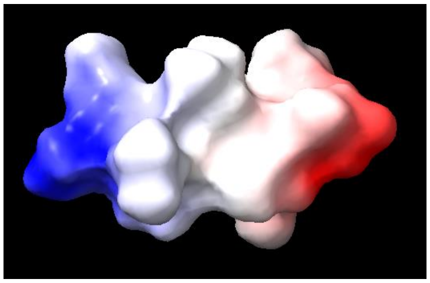 |

**Pair 38** **LGAVQNEV** **LTAVAEEV**

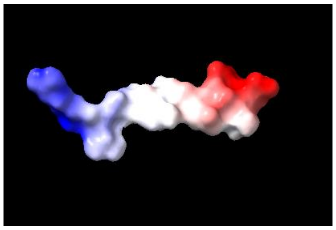

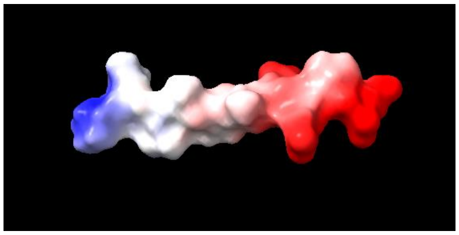

**Pair 39** **LYVLFEV** **LYHVYEV**

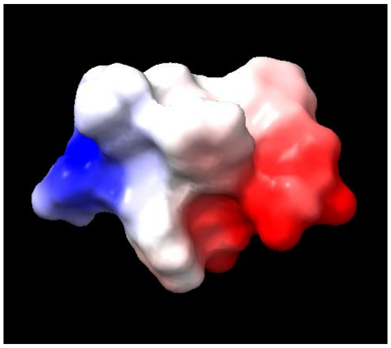

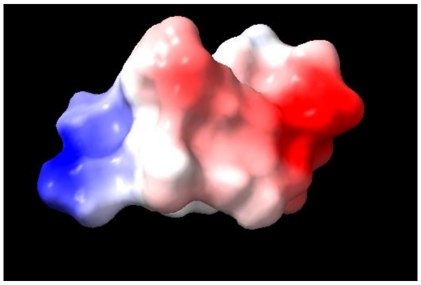

**Pair 7** **AVNGVLW** **AVDGVLS**

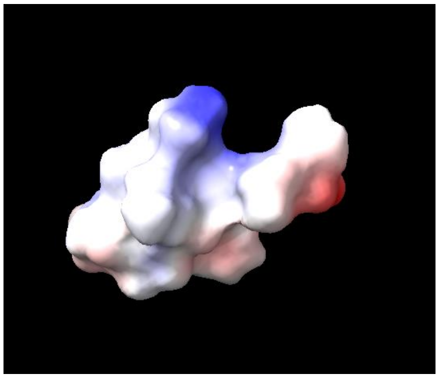

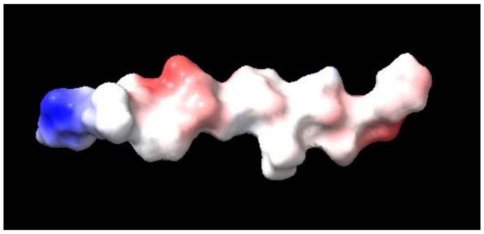
