## Supplementary material for "Structural Homology and Electrostatic Potential Comparisons of Epitope Pair Candidates for Molecular Mimicry Triggering of Type 1 Diabetes Mellitus": All suplementary material: Supplementary material.docx

**Supplementary Materials**

**Table ST1: C**omplete results of the work presented here. Pairs are listed in alphabetical order by the E_T1D_ epitope first aminoacidic. Amino acid letters in RED correspond to the original sequences of the epitopes, while those in back are those added to satisfy the minimum length needed for AlphaFold predictions. The color in the Antigen fold column is associated with the sequences depicted in the same color when comparing structure of isolated epitopes to the one in the full antigen in Figs. 2-10.

**Table ST2**: RMSD Between Overlapping E_INF_/E_T1D_ pairs when sequences are pruned to the best overlapping sequence. Amino acids in red correspond to the original epitopes, those in black are added to reach the minimum required by AlphaFold.

**Figure S1:** Comparison of the calculated electrostatic potentials for all the nonmatching pairs from Table 3. The palette options were colored as follows: blue is defined to be negative; red is defined to be positive, and white is defined to be neutral. The nomenclature of the pair numbers can be found in Table ST1 of the supplementary material.

**Figure S2:** Comparison of the calculated electrostatic potentials for the epitope pairs showing discrepancies in the folding secondary structure in Tables 2-10. The palette options were colored as follows: blue is defined to be negative; red is defined to be positive, and white is defined to be neutral. The nomenclature of the pair numbers can be found in Table ST1 of the supplementary material.
