## Supplementary material for "Structural Homology and Electrostatic Potential Comparisons of Epitope Pair Candidates for Molecular Mimicry Triggering of Type 1 Diabetes Mellitus": All suplementary material: Table ST2.docx

**Table ST2**: RMSD Between Overlapping E_INF_/E_T1D_ pairs when sequences are pruned to the best overlapping sequence. Amino acids in red correspond to the original epitopes, those in black are added to reach the minimum required by AlphaFold.

| **E-T1D (EXT)** | **E-INF (EXT)** | **Pruned RMSD (Å)** | **Number of Residues** |
| --- | --- | --- | --- |
| HLLIDLTSFLL | PLLYDANYFLC | 0.09 | 3 |
| LPLLALLALWG | LLLEWLAMAV | 0.14 | 9 |
| PKTRREAEDL | FLTRVEAQLH | 0.26 | 5 |
| LLPLLALLALW | GAFLLGLLFFV | 0.28 | 10 |
| LPLLALLALWG | VFLLVTLAILT | 0.33 | 11 |
| LPLLALLALWG | LTLLSTAALPV | 0.34 | 9 |
| LLPLLALLAL | LDHLLVLLEK | 0.39 | 10 |
| ASLYHVYEVNL | TLLYVLFEVFD | 0.44 | 8 |
| PLLALLALWGP | TGEALWALPHA | 0.45 | 3 |
| WMRLLPLLAL | LDHLLVLLEK | 0.46 | 10 |
| PKTRREAEDLQ | EARRRLAEMCF | 0.48 | 9 |
| LPLLALLALW | LLLLTLLATV | 0.5 | 10 |
| LIDVFHQYSG | YADVFHLYLQ | 0.55 | 10 |
| FALTAVAEEV | KVLTLFAEVE | 0.56 | 10 |
| YALLGIPLNVI | MALLGLTLGVL | 0.58 | 4 |
| CGSHLVEALYL | TMADLVYALRH | 0.61 | 10 |
| GSHLVEALYL | MADLVYALRH | 0.69 | 10 |
| GYIVTDQKPLS | GYIPLVGAPLG | 0.69 | 6 |
| LAEYLYNIIKN | PALYLYNTGRS | 0.73 | 11 |
| TILKDFSILLM | SILSPFLPLLP | 0.74 | 8 |
| MRLLPLLALLA | LLLEWLAMAV | 0.81 | 9 |
| MLMPVHFLL | KLMPVCVET | 0.84 | 7 |
| LAVDGVLSVHS | IFVDGVPFVVS | 0.85 | 7 |
| PLLALLALWGP | REHALLAFTLG | 0.86 | 9 |
| AMVALIDVFHQ | EKMALYDVVSK | 0.9 | 11 |
| GLQANPVEVQS | IMDPNPPEVVS | 0.93 | 7 |
| GLLPLLALLA | QWLPTGTLLV | 0.96 | 3 |
| PLTAVAEEVE | RLGAVQNEVI | 0.99 | 10 |
| GYIVTDQKPL | SMIVTDHRYV | 1.04 | 10 |
| RLLPLLALLA | LLLCLIFLLV | 1.04 | 10 |
| ILAVDGVLSVH | ATAVNGVLWTV | 1.06 | 4 |
| WMRLLPLLALL | GAFLLGLLFFV | 1.07 | 11 |
| RLLPLLALLA | FLLVTLAILT | 1.08 | 10 |
| MRLLPLLALL | LLLLTLLATV | 1.09 | 8 |
| GGLQANPVEVQ | KALQFIPVESL | 1.09 | 9 |
| LALWGPDPAAA | EALWALPHAAA | 1.1 | 5 |
| ILAVDGVLSV | FLPVDFFPSV | 1.16 | 10 |
| IVTDQKPLSLAA | IVSTAAQTFLAT | 1.27 | 8 |
| IVTDQKPLSLAA | IVVNASASEAAA | 1.31 | 9 |
| LALWGPDPAAA | KLLPDTAAV | 1.35 | 7 |
| PLLALLALWG | KYLAAYALVG | 1.46 | 3 |
| WMRLLPLLAL | LSPFLPLLPI | 1.67 | 4 |
| PRLLPLLALLE | LRLNELLAYVS | 96 | 11 |
